# Effects of high dosage methamphetamine on glutamatergic neurotransmission in the nucleus accumbens and prefrontal cortex

**DOI:** 10.1101/2021.04.22.440987

**Authors:** Devesh Mishra, Jose I Pena-Bravo, Shannon M Ghee, Carole Berini, Carmela M Reichel, Antonieta Lavin

## Abstract

**Rationale:** Methamphetamine (METH) induces changes in the glutamatergic system and elicits cellular alterations in the cortico-accumbens circuit.

**Objective:** While there is a body of literature on the effects of METH on dopaminergic transmission, there is a gap in knowledge regarding the effects of a high dose of METH on synaptic glutamatergic neurotransmission, specifically in brain regions involved in goal directed behavior (nucleus accumbens core; NAc core) and executive functions (prefrontal cortex;PFC).

**Methods:** In order to fill that gap we assessed synaptic glutamatergic transmission using a well established METH administration regime (4 × 4 mg/kg ip at 2 hr intervals) followed by 7 days of abstinence. Rats were then sacrificed and whole cell and field recordings were performed in the NAc core and medial PFC.

**Results:** METH treatment elicited a significant decrease in paired pulse ratio in NAc core and a significant increase in AMPA/NMDA ratio driven by increases in AMPA currents. On the other hand, there were no significant changes in measures of synaptic glutamate in the PFC.

**Conclusion:** These results suggest that a high dose of METH treatment followed by a period of abstinence elicits significant increases in indices of glutamatergic transmission in the NAc core with no detectable changes in mPFC, denoting that neurons and glutamate terminals in this limbic region have a higher susceptibility to a neurotoxic METH regime.

## Introduction

Methamphetamine (METH) is one of the most abused and highly potent psychostimulants available and its use has increased exponentially in the last decade (United Nations Office on Drug and Crime report, 2014). While there are a number of studies on the effects of METH in striatal regions, less is known regarding the synaptic effects of a high dose of METH in the cortico-accumbens circuit. The medial prefrontal cortex (mPFC) and nucleus accumbens (NAc) are essential components of the addiction circuit (Robinson and Kolb, 1997) and are involved in regulating executive function and goal directed behavior respectively, thus, using a high METH dose protocol, we investigated the effects of the psychostimulant in synaptic glutamatergic transmission in the mPFC-NAc core circuit.

The majority of the literature dealing with the cellular effects of METH in the cortico-accumbens circuit is focused in drug self -administration or sensitization protocols. Thus, it is not surprising that different and sometimes contradictory changes are reported. Brady and colleagues (2003) reported that a METH-sensitization regime (5.0 mg/kg /5 days) increases the duration of up states and decreases excitability in NAc neurons recorded *in vivo*. Similarly, Graves and colleagues (2015) show that METH self-administration (0.1 mg/kg/0.1 ml for 14 days) decreased the excitability of NAc shell neurons. In the PFC, an acute METH bolus in rats (30 mg/kg, ip), decreased the levels of NR1A NMDA receptor subunits and increased levels of NR2A and GluA2 in the PFC without changes in striatum (Simoes et al 2008). Furthermore, Herrold and colleagues (2013) report that following a METH sensitization regime (1 mg/ml for 2 days) in rats, levels of GluA2 surface expression in the cortex were increased. We have shown that long access METH self administration (0.2 mg/infusion/6 hours/14 days) increased *in vivo* basal firing frequency accompanied by an increase in the number of bursting cells in the PFC (Parsegian et al., 2013). Recently, Lominac and colleagues (2016) using a repeated low dose METH regime (2 mg/kg/10 injections) show that the regime induce glutamate release and lower indices of NMDA receptor expression in C57BL/6J mice, but did not alter basal extracellular glutamate content.

In humans, METH induces striatal neurotoxicity indexed as a decrease in dopamine transporters, which is indicative of dopamine terminal degradation (McCann et al., 1998; Sekine et al., 2006). METH neurotoxicity has also been established pre-clinically in rodents consistently demonstrating that high dosages of non-contingent METH cause a loss of striatal dopamine transporters (Wagner et al., 1980; Eisch et al., 1992), decreased tyrosine hydroxylase (Knapp et al., 1974; Broening et al., 1997), and increased GFAP levels (Pu and Vorhees, 1993; LaVoie et al., 2004). Further, acute METH results in marked increase in locomotor activity (Kubota et al., 2002) and high dose regimens induce stereotyped behavior, which can be indicative of psychosis (Abekawa et al., 2008). Co-existing with these acute physiological effects, METH (4 × 4 mg/kg, 2 hr intervals) also causes memory impairments at least one week following treatment (Schroder et al., 2003; Belcher et al., 2005; Reichel et al., 2012; Heysieattalab et al., 2015).

Several studies have shown that exposure to high doses of METH dysregulates the dopaminergic system in the dorsal striatum, however the ventral striatum (i.e., nucleus accumbens) appears to be less affected by the drug adverse effects (for a review see Kasnova and Cadet, 2009; but also see Kuhn et al., 2011). There is a dearth of reports regarding effects of high METH dosage on excitatory neurotransmission in the NAC or PFC. Here we report that a high dosage METH regime (4 × 4 mg/kg, 2 hrs intervals) elicited an increase in glutamate indices in the NAc core. In contrast, no significant changes in glutamatergic synaptic transmission were found in the mPFC.

## Methods

Sprague Dawley male rats received four ip injections of 4 mg/kg methamphetamine hydrocholoride (dissolved in 0.9% saline Sigma, St Louis, MO, USA) every two hours for 8 hours with rectal probe body temperature measurements every hour. Following this neurotoxic regime, rats were individually housed with free access to food and water for 7 days, on 12 hour reversed light/dark cycle. All procedures were conducted in accordance with the ‘Guide for the Care and Use of Laboratory Rats’ (Institute of Laboratory Animal Resources on Life Sciences, National Research Council) and approved by the IACUC of the Medical University of South Carolina.

On day 8, rats were anesthetized with isoflurane (Abbott Laboratories) and the brain was removed. Coronal slices containing medial PFC (300μm) or NAc core (230 or 400μm) were cut in ice-cold high-sucrose solution containing (in mM): sucrose, 200; KCl, 1.9; Na_2_HPO_4_, 1.2; NaHCO_3_, 33; MgCl_2_, 6; CaCl_2_, 0.5; D-Glucose, 10; ascorbic acid, 0.4 using a Leica VT 1200 S vibratome. Slices were incubated at 31°C for at least 1 h before recordings; the incubation medium contained (in mM): NaCl, 125; KCl, 2.5; NaH_2_PO_4_,1.25; NaHCO_3_, 25; MgCl_2_, 4; CaCl_2_, 1; D-Glucose, 10; sucrose, 15; ascorbic acid, 0.4, aerated with 5% CO_2_/ 95%O_2_. Following incubation, slices were transferred to a submerged chamber and superfused with oxygenated artificial cerebrospinal fluid (aCSF) (in mM): NaCl, 125; KCl, 2.5; Na_2_HPO_4_, 1.2; NaHCO_3_, 25; MgCl_2_, 1.3; CaCl_2_, 2.0; D-Glucose, 10 and ascorbic acid, 0.4 at room temperature.

### Field potential or population spikes (PS) recording

A glass recording electrode filled with rACSF (1-2 MΩ resistance in situ) was placed in close proximity to a concentric bipolar stimulation electrode. An input/output response curve was generated by stepwise increases of the stimulation intensity from 10 μA to 100 μA at 4 Hz. The stimulation pulse had a duration of 0.1 msec. Stimulation intensity resulting in 50 percent of the maximum output response was used for baseline recordings. For paired-pulse experiments, the inter stimulus interval was set to 50 ms stimulating at 0.033 Hz. Recordings were performed for 30 minutes and the last 5 minutes of recording were averaged for analysis and statistical comparison. Recordings were made using a Axopatch 200B amplifier (Axon Instruments, CA), connected to a computer running Windows XP and Axograph X software. Signals were amplified 500 times, acquired at 10 kHz and filtered at 2 KHz.

### Voltage clamp recordings

*Voltage clamp recordings*: Picrotoxin (50 μM) was included in the perfusion solution to block GABA_A_ receptors. For voltage-clamp recordings, electrodes (2.5-3.5 MΩ resistance in situ) were filled with a solution containing (in mM): CsCl, 135; HEPES, 10; MgCl2, 2; EGTA, 1; NaCl, 4; Na-ATP, 2; tris-GTP, 0.3; QX-314, 1; phosphocreatine, 10; 285 mOsmols. Series resistance (10-25 MΩ) was continually monitored throughout the experiment via a −5 mV (50 ms) hyperpolarizing pulse. Neurons were clamped at −70 mV and spontaneous excitatory postsynaptic currents (sEPSCs) and paired pulse responses were recorded. For AMPA/NMDA experiments, EPSC’s were evoked using a bipolar concentric electrode placed within 300-400 μm of the recording electrode. Neurons were clamped at +40 mV and evoked currents were recorded for 5 min at 0.033 Hz, then the NMDAR antagonist AP5 (D(-)-2-amino-5-phosphonovaleric acid (50 mM) was applied to isolate AMPA currents. Following 5 minutes of D-AP5 perfusion, eEPSC’s were recorded for an additional 5 minutes. The NMDA currents were obtained by digital subtraction (I_Total_− I_AMPA_) and the AMPA to NMDA ratio was calculated with the following formula: I_AMPA_ / I_Total_ – I_AMPA_. Paired pulse experiments involved two evoked EPSCs with an interval inter stimulus of 50 msec at 0.033Hz. Recordings were made using a Multiclamp 700B amplifier (Axon Instruments, CA), connected to a computer running Windows XP and Axograph X software. All recordings were obtained from pyramidal neurons in layers V/VI of the prelimbic cortex, identified using infrared-differential interference contrast optics and video-microscopy.

## Data analysis

Data was analyzed using unpaired 2-tailed t-tests or analyses of variance (ANOVAs) followed by Bonferroni’s post-hoc test**p<0.01 when appropriate.

## Results

Using field and whole cell recordings we assessed sEPSC frequency and amplitude, AMPA/NMDA ratio and paired pulse responses in brain slices containing the NAc core or mPFC of animals treated with high doses of METH (n=7 animals) or saline (n=5 animals). There were no differences in body weight between METH and saline animals over the course of the injections, nor did METH increase body temperature relative to baseline (data not shown). We have previously shown that this METH regime produces marked neurotoxic consequences in our laboratory (Reichel et al., 2012). In that study we did not observe changes in body temperature but still report reductions in dopamine transporters and tyrosine hydroxylase and elevated GFAP levels in the dorsal striatum.

### NAc core

Population spike recordings in the NAc core revealed a significant depression of paired-pulse ratio (PPR) following METH (Fig. 1b, t(7)=3.01 p<0.01). The average paired pulse ratio in saline controls was 0.97 ± 0.03 versus 0.86 ± 0.1 in METH-treated rats, suggesting an increase in neurotransmitter release probability in METH-treated rats. A two way ANOVA revealed main effects of stimulus intensity (F[9, 190]=47.52, p<0.0001) and treatment group (F[1,190)=47.1, p<0.0001) but no interaction. The mean values of the input/output (I/O) curve show a significant decrease in synaptic output in METH-treated rats compared to saline controls (p< 0.0001 Fig 1c).

**Figure 1.**
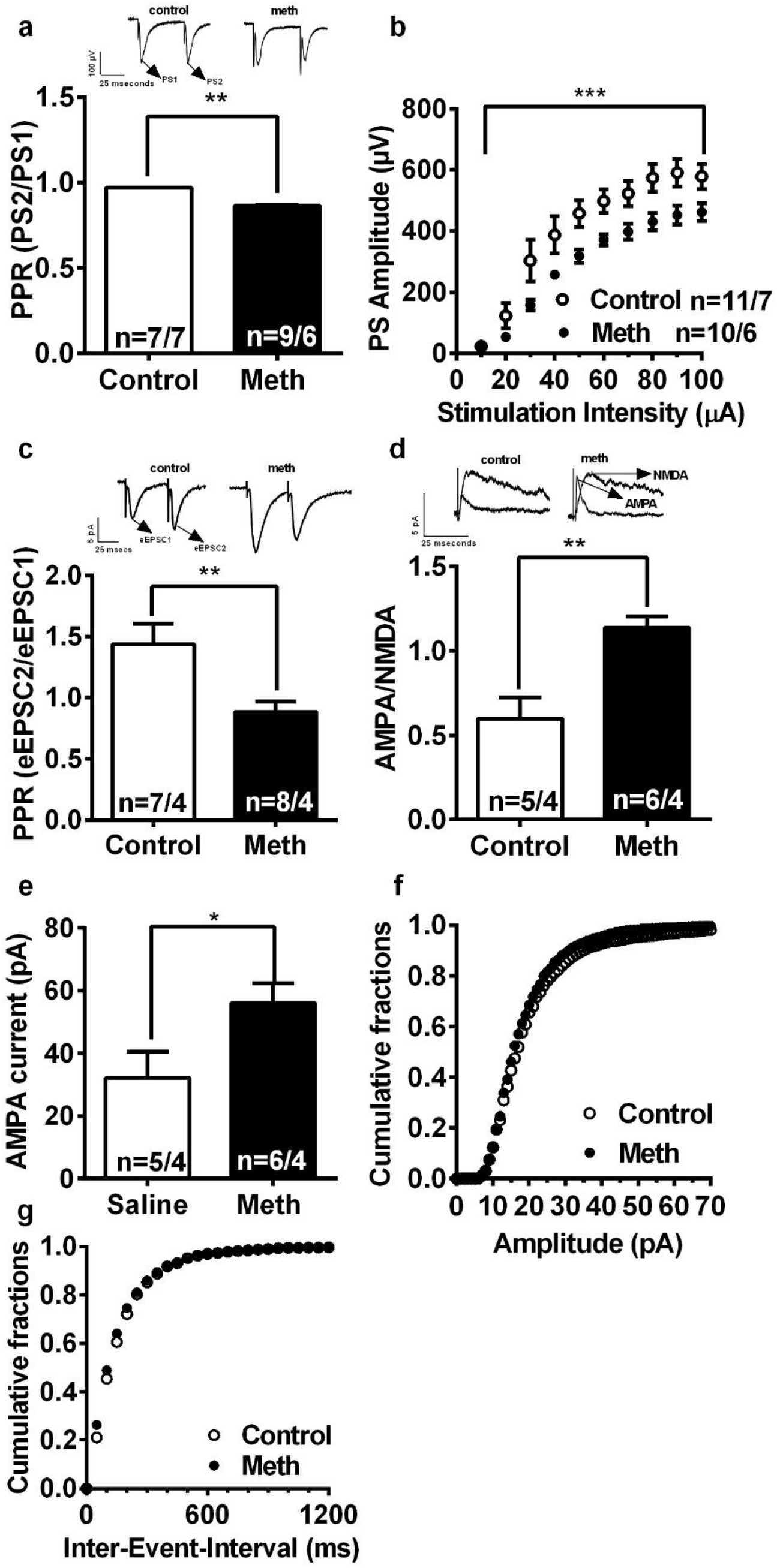
Effects of METH toxic regime on synaptic activity in the NAc. a) Representative traces of paired pulse ratio (PPR) obtained with field recordings in the NAc. b) PPR show a significant depression in METH-treated rats, (**t(7)=3.01 p<0.01) when compared with control animals, suggesting an increase in the probability of release of glutamate. n=number of fields recordings/number of rats. c) In METH-treated rats, the I/O curve obtained with field recordings shows a significant right shift (*** p<0.0001) compared to control animals, indicating a significant decrease in synaptic output in METH-treated rats. d) Representative traces of PPR obtained with whole cell recordings in NAc neurons. e) Whole cell recordings of NAc neurons show a significant decrease in PPR in METH animals when compared with controls (**t(13)=3.08, p<0.01) suggesting an increase in the probability of release of glutamate. n= number of cells/number of rats. f) Representative traces of AMPA/NMDA ratio for control and METH animals. g) AMPA/NMDA ratio shows that METH toxic regime elicits a significant increase in the ratio (**t(9)=3.98, p<0.01), indicative of a postsynaptic potentiated state. e) Comparing mean evoked AMPA currents shows that METH rats exhibit a significant increase when compared to control rats (control rats: 32.1 ± 8.3 pA versus 56.0 ± 6.2 pA in METH-treated rats; paired t-test, two tail * p< 0.04).

Voltage clamp recordings from medium spiny neurons identified via infrared-differential interference contrast optics and video-microscopy, also showed a significant depression of PPR in METH-treated rats when compared to controls (Fig. 1e, t(13)=3.08, p<0.01).The average paired pulse ratio of saline control rats was 1.44 ± 0.16 vs 0.88 ± 0.08 in METH-treated rats. Furthermore, we observed a significant increase in AMPA/NMDA ratio in METH-treated rats relative to controls (Fig. 1g, t(9)=3.98, p<0.01), which suggest a postsynaptic potentiated state.

When individual synaptic currents (AMPA and NMDA) were isolated, it was found that AMPA currents exhibit a significant increase in METH rats when compared to saline rats (saline rats: 32.1 ± 8.3 pA versus 56.0 ± 6.2 pA in METH-treated rats; paired t-test, two tail p< 0.04, Fig 1h). No changes were observed upon comparing sEPSCs averages of amplitudes or inter-event-intervals.

In summary, a neurotoxic METH dosage elicits an increase in glutamate synaptic indices in the NAc core as well as an increase in the AMPA/NMDA ratio mediated by significant increases in AMPA currents. In contrast, we found a significant shift to the right in the I/O curve, suggesting a decrease in NAc medium spiny neurons intrinsic excitability.

### mPFC

In mPFC, whole-cell recordings from deep layer pyramidal neurons identified using infrared-differential interference contrast optics and video-microscopy, revealed no changes in PPR (Fig 2b) or AMPA/NMDA ratio (Fig. 2d) in METH-treated rats compared to controls. No differences were observed upon comparing cumulative distributions of sEPSCs inter-event-intervals or average frequencies. Since no changes were found in the PFC using voltage clamp, we did not record any population spikes in this region. Interestingly, our results suggest that a single day high dosage METH regime does not affect glutamatergic synaptic indices in the mPFC.

**Figure 2.**
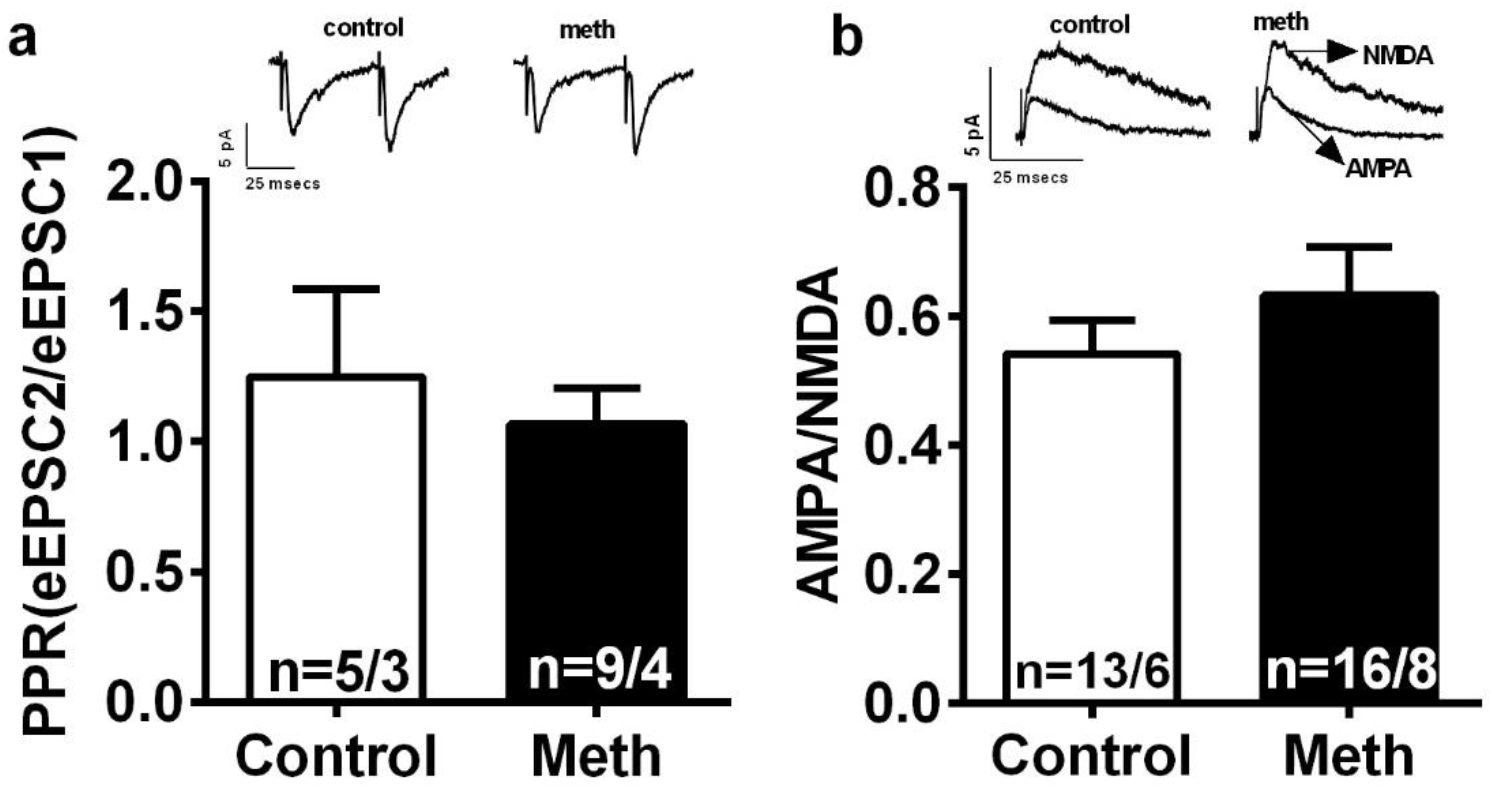
Effects of METH toxic regime in synaptic activity in the PFC. a) Representative traces of paired pulse ratio (PPR) obtained with whole cell voltage clamp. b) Whole cell voltage clamp PPR recordings shows that METH toxic regime does not affect probability of release of glutamate in the PFC. c) Representative traces of AMPA/NMDA ration in control and METH animals recorded in the PFC. d) METH toxic regime does not affect the AMPA/NMDA ratio in the PFC. n=number of cells/ number or rats.

## Discussion

World-wide METH use is increasing at a rapid rate; therefore, it has become increasingly important to understand the occurring changes in synaptic physiology following METH exposure. Given the relevance of goal directed behavior and executive function in addiction, we investigated pre- and post- synaptic changes in the cortico-accumbens pathway following acute METH exposure. Corticostriatal glutamate neurotransmission mediates addictive behavior, and synaptic changes in this circuit are relevant for understanding mechanisms mediating addiction. Here we show that a high dosage METH regimen increases glutamate release in the NAc core but, surprisingly does not affect glutamatergic transmission in the medial PFC.

We found that in the NAc core, METH treatment produced paired pulse depression (PPD) in glutamatergic synapses, suggesting increased synaptic glutamate release (Mennerick and Zorumski, 1995; Debanne et al., 1996; Bonci and Williams, 1997; Regehr 2012). PPD is a presynaptically mediated phenomenon and it has been suggested that the increase in neurotransmitter release results from a depletion of the ready releasable pool of vesicles following the first stimuli (Rosemund and Stevens, 1996; Wang and Kaczmarek, 1998; Oleskevich et al., 2000; Meyer et al., 2001), however other mechanisms such as presynaptic calcium channel inactivation (Gingrich and Bryne, 1985), negative feedback trough inhibitory autoreceptors (Forsythe and Clements 1990) or receptor desensitization (Otis et al., 1996; Rozov et al., 2001) may be mediating the increase in glutamate transmission. Our finding of an increase in synaptic glutamate index is complemented by the observed increase in AMPA/NMDA ratio.

In contrast, we also observed a right shift in the NAc core I/O curve and a decrease in the maximum response, suggesting a decrease in synaptic excitation or decrease in cell viability/terminal number. The increase in indices of synaptic glutamate and the shift in the I/O curve may seem contradictory. We reconcile these results by proposing that the increase in synaptic glutamate indices is mediated via a presynaptic mechanism (see above) and the right shift in the I/O curve is mediated via a postsynaptic mechanism, i.e., a decrease in intrinsic neuronal excitability of the NAc core cells. Indeed, it has been shown that I/O curves are the result of postsynaptically-mediated AMPA currents (Mishra et al., 2013; Morud et al., 2015) and Brady and colleagues (2003) as well as Graves et al (2015) have show that NAc core cells from METH treated rats exhibited a decrease in excitability.

Psychostimulants have been repeatedly shown to affect AMPA to NMDA function in reward related brain regions (Kourrich et al., 2009; Murata et al., 2009; Moussawi et al., 2009; Mu et al., 2010; Zhang et al., 2002), these changes are thought to be the driving force behind the idea of drugs of abuse ‘hijacking’ the reward circuitry (Moussawi et al., 2009). Upon isolating individual synaptic currents mediated by AMPA and NMDA, we found increased synaptic AMPA currents in METH treated rats. This increased AMPA function has been reported earlier following psychostimulant exposure and is directly associated with AMPA receptors mediating drug seeking (Bowers et al., 2010; Ferrario et al., 2011; Millan and McNally, 2011; Pierce and Wolf M, 2013; Loweth et al., 2014; Chen and Chen 2015; Terrier et al., 2015; Jedynak et al., 2016). AMPA infusions into the NAc core reinstate drug seeking whereas infusion of AMPAR antagonists blocks reinstatement produced by systemic cocaine, NAc core DA, or cocaine infusions in the prefrontal cortex (Cornish et al., 1999; Cornish and Kalivas, 2000; Park et al., 2002). Overall, the activation of AMPA receptors in the NAc core via glutamatergic afferents from mPFC seems to mediate relapse to drugs of abuse. Hence these changes in pre- and post-synaptic glutamate neurotransmission, even if transient, do suggest that METH along with its general psychostimulant effects, increase glutamate transmission in these addiction-associated brain regions. However, Herrold and colleagues (2013) and Nelson et al, (2009) have reported no change in AMPAR in the NAc of METH or amphetamine rats. There are obvious differences between our methods and those used by Herrold and Nelson. The most obvious one is the difference in METH dosage: we use a high dosage METH protocol and Herrold and colleagues used either a single acute injection of 1 mg/kg of METH or a repeated treatment using the same dose for 3 days followed by 14 day withdrawal. Nelson et al used either a single 2.5 mg/kg amphetamine injection or a sensitization regime with different withdrawals times, thus comparing results it is not feasible.

On the other hand, our results show no changes in synaptic glutamate transmission in the mPFC. This lack of modifications in glutamatergic synaptic transmission may reflect a recovery to basal levels after 7 days abstinence. Alternatively, this may suggest that glutamatergic inputs to PFC are not affected by the METH regimen used in our experiments due to the passive drug delivery system. Since we did not find any changes in cortical glutamate, our results may suggest that other non-cortical glutamatergic inputs are mediating the changes observed in the NAc, however, those experiments were out of the scope of the present manuscript. Lastly, our results may suggest that NAc core glutamatergic afferents are particularly sensitive to METH neurotoxic regimes.

## Conclusion

Our results suggest that high doses of METH alter the NAc core, which is involved in goal directed functions, without affecting the PFC that mediates mainly executive functions.

## Abbreviations

Nucleus Acumbens: NAc
Prefrontal Cortex: PFC
Methamphetamine: METH
PPR: Paired Pulse Ratio

## Acknowledgements

This research was supported by National Institute on Drug Abuse (NIDA) grants DA033049 (CMR), DA007288 (JPB), and NIH grant C06 RR015455.

## Notes

The authors do not have any conflict of interest to disclose

### Competing Interest Statement

The authors have declared no competing interest.

